# panHiTE: a comprehensive and accurate pipeline for TE detection in large-scale population genomes

**DOI:** 10.1101/2025.02.15.638472

**Authors:** Kang Hu, Minghua Xu, Jianxin Wang

**Affiliations:** School of Computer Science and Engineering, Central South University, Changsha, 410083, China; Xiangjiang Laboratory, Changsha, 410205, China; Hunan Provincial Key Lab on Bioinformatics, Central South University, Changsha, 410083, China

**Keywords:** TE detection, genome annotation, LTR-RT, large-scale population, Nextflow pipeline

## Abstract

Transposable elements (TEs) are key drivers of genomic variation and species evolution. Advances in high-throughput sequencing have enabled whole-genome sequencing of individuals or subspecies, facilitating the identification of population-specific variations. Detecting population-specific TE insertions at scale is crucial for understanding species-specific phenotypic traits. However, tools for constructing comprehensive pan-TE databases remain limited. To address this gap, we develop panHiTE, a population-scale TE detection and annotation tool with several core innovations. panHiTE features a deep learning-based long terminal repeat retrotransposon (LTR-RT) detection algorithm, outperforming existing tools in both sensitivity and precision. It also introduces a novel de-redundancy algorithm, which eliminates highly divergent redundant TE instances, significantly reducing the size of the TE library. Additionally, panHiTE can detect low-copy TEs, which are overlooked in individual genome analyses and absent from existing databases due to their rarity. Furthermore, panHiTE allows for TE-gene association analysis, enabling comprehensive insights into TE-driven phenotypic variation. panHiTE, powered by a Nextflow pipeline, enables efficient and scalable TE detection in large plant genomes and has successfully been applied to hundreds of plant population genomes, demonstrating its effectiveness and scalability.

## Introduction

Transposable elements (TEs) are ubiquitous components of genomes and serve as one of the primary drivers of genome evolution^1,2^. They contribute to host genome evolution through processes such as chromosomal rearrangements, exon shuffling, and the donation of coding sequences^3–5^. TEs can directly alter transcript structures by inserting into genes^6^ or act as “ready-to-use” cis-regulatory elements by providing abundant transcription factor binding sites when inserted upstream or downstream of genes^7,8^. TEs are also implicated in human diseases^9,10^ and plant phenotypes^11–13^. For instance, LINE-1 insertions disrupt tumor suppressor genes, driving mutations in colorectal cancer^14^. In plants, Barbara McClintock first demonstrated the role of TEs in phenotypic variation by linking them to color variegation in maize kernels and leaves in 1948^15^. Similarly, LTR retrotransposons have been associated with key traits, such as red skin in apples due to an insertion upstream of the MdMYB1 gene^16^, loss of fruit color in grapes caused by the Gret1 LTR retrotransposon^17^, and enhanced apical dominance in maize through a TE insertion in the regulatory region of the tb1 gene^18^.

Despite their importance, comprehensive studies of TEs face significant challenges. Plant genomes are often large, highly repetitive, and structurally complex, making it difficult to fully characterize their TE content using traditional genome annotation methods^19–22^. Large-scale TE detection in plant pan-genomes is further hindered by long processing times, incomplete results, and high false-positive rates^23,24^. Additionally, the assembly of repetitive regions^25,26^, the limited availability of genome assemblies within species, and the complexity of annotating diverse TE classes^27^ pose further obstacles. Current TE detection tools suffer from issues such as high false-positive rates^28,29^, excessive fragmentation^30,31^, and limited scalability, particularly when applied to large and complex plant genomes^32^. At the same time, incomplete or outdated TE databases prevent a comprehensive understanding of TE diversity and the identification of population-specific TE insertions^33,34^.

Recent advances in long-read sequencing and improved assembly algorithms have provided new opportunities for studying population-level variation in TEs^35,36^. These technologies allow for the construction of multiple genome assemblies within species, enabling a more nuanced exploration of genome evolution and helping enhance crop breeding^37–39^. However, these advances also bring new challenges in terms of data processing, the integration of large datasets, and the accurate detection of rare or low-copy TEs. There is an urgent need for a scalable, accurate, and efficient pan-TE detection tool that can handle the complexities of large genome datasets and provide a more complete picture of TE dynamics across populations.

To address these challenges, we have developed panHiTE, an enhanced version of the TE annotator HiTE^40^, designed to significantly improve the accuracy and scalability of TE detection in pan-genomes. One of the key innovations of panHiTE is the incorporation of FiLTR, a novel LTR retrotransposon detection tool developed in this study, which significantly boosts detection accuracy for LTR retrotransposons. Furthermore, panHiTE includes structural and domain validation specifically tailored for low-copy candidate TEs, ensuring accurate identification of even rare or poorly represented families. A major feature of panHiTE is its progressive redundancy removal algorithm, which effectively eliminates divergent redundant TE instances, thereby reducing the size of the pan-TE library and improving both computational efficiency and result accuracy. Additionally, panHiTE leverages population-based pan-genomes to recover low-copy TEs, identifying their copies across multiple ecotypes and addressing the challenge of detecting TEs that are often missed in individual genome analyses. Extensive testing has demonstrated that panHiTE outperforms other commonly used TE detection tools, such as panEDTA^41^, in both sensitivity and precision. Built on the Nextflow workflow^42^, panHiTE is capable of parallel processing on high-performance computing (HPC) platforms, enabling efficient large-scale TE detection and annotation. The tool also features checkpoint recovery, ensuring robustness in long-running computational tasks. Notably, panHiTE has been successfully applied to annotate TEs in 17 wheat genomes^43^ (14 GB each), providing a powerful solution for TE analysis in ultra-large pan-genomes. Overall, panHiTE offers a significant advancement in the study of TE evolution and its contribution to phenotypic diversity in plants.

## Results

### The workflow of panHiTE

With the advancement of sequencing technologies, we are able to obtain DNA and RNA sequencing data from a large number of individuals across different ecotypes (Fig. 1a). By utilizing population-level data, fast and accurate TE detection and annotation can help researchers identify TE insertions that contribute to individual-specific traits. As shown in Fig. 1b, panHiTE begins by performing TE detection on the individual genomes within the population using HiTE. Given the high redundancy of TEs across genomes, panHiTE leverages the progressive redundancy removal algorithm to eliminate redundant TE instances. Low-copy TEs detected in individual genomes are re-aligned to the pan-genomes to obtain sufficient copy numbers, allowing the recovery of the true TEs. Reliable TEs are merged and processed through the de-redundancy module to generate a pan-TE library. This library is then used to annotate each genome, producing corresponding TE annotations. Following this, panHiTE conducts association analysis by examining TE and gene annotations to identify TEs inserted upstream, inside, or downstream of genes, revealing structural variations caused by TE insertions across the population (Fig. 1b). Finally, panHiTE uses RNA-seq data to quantify both TEs and genes, enabling the identification of TEs with significant expression differences at the population level, as well as TE insertions that cause differential gene expression, referred to as “TE-induced differential expression loci” (TIDELs). These specific TE insertions can affect gene transcripts and protein production, ultimately influencing individual phenotypes (Fig. 1c).

**Fig. 1.**
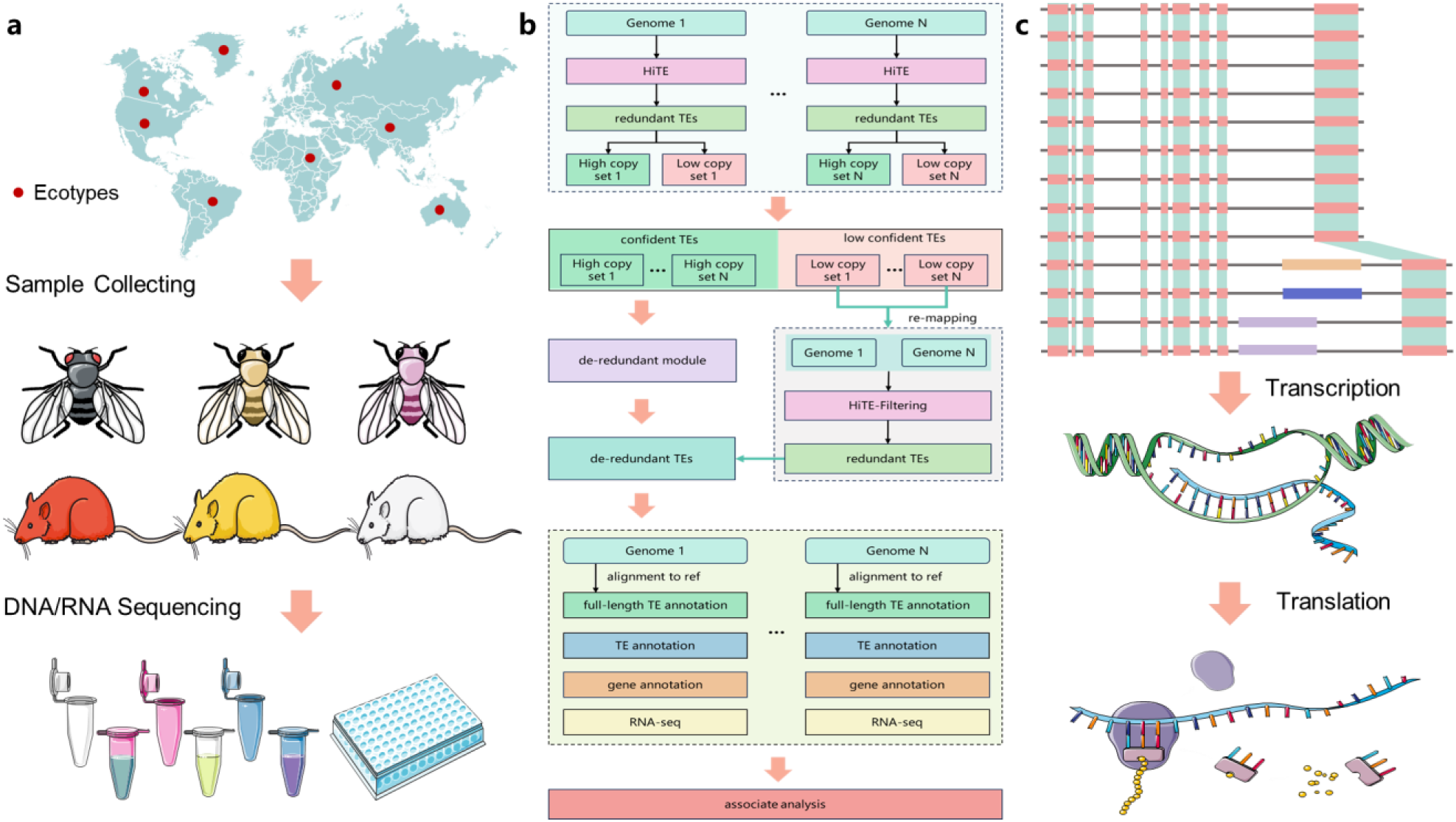
Pan-genome annotation and analysis of different accessions using panHiTE. **a.** A broad range of biological specimens from geographically distinct regions are acquired to perform DNA and RNA sequencing analyses. **b.** The panHiTE workflow. HiTE is used to detect TEs in each genome, and TEs with low copy numbers are collected and re-mapped to the pan-genomes to obtain additional copies. All TEs undergo a de-redundant module to remove redundant and are subsequently used for pan-genome annotation. TE and gene annotations, combined with RNA-seq reads, are used for association analysis to detect population-based structural variations that may lead to changes in gene expression. **c.** Different types of LTR-RT insertions within the gene in specific ecotypes generate diverse transcript variants, potentially altering protein structure and expression levels.

Accurately identifying the boundaries and structure of a TE relies on the presence of multiple copies. However, when a TE exists in low copy numbers within the genome, determining its true boundaries and structure becomes challenging. This limitation frequently results in the omission of such low-copy TEs from analyses. Leveraging the pan-genomes, panHiTE realigns low-copy TEs detected in individual genomes across multiple genomes, thereby obtaining additional copies and helping to recover some of the true low-copy TEs. panHiTE has been evaluated on 32 *A. thaliana* accessions, identifying 172 candidate low-copy TE insertions. After removing redundancies and excluding TEs overlapping with those in the detected pan-TE library, 41 unique TE families remained, including 38 TIR and 3 Helitron families. By querying the Repbase database and manual inspection, 4 TIR and 1 Helitron families are found to be false positives. Additionally, 21 families are shared between panHiTE and Repbase, while the remaining 15 TIR families are confirmed as genuine through manual validation, representing novel transposons identified by panHiTE that are not yet recorded in Repbase. This demonstrates that panHiTE can accurately recover missed low-copy TEs using the pan-genomes.

Compared to existing tools, panHiTE is capable of identifying more full-length TEs, reducing the need for manual editing. With higher precision and sensitivity, panHiTE improves the quality and completeness of TE annotation. By automating the integration of TEs, genes, and RNA-seq reads, panHiTE identifies population-specific TE insertions and differentially expressed genes caused by these insertions, which may serve as key drivers of species evolution. Built on the Nextflow workflow, panHiTE enables automated multi-node parallel execution on HPC clusters, making it highly suitable for the annotation and analysis of pan-genomes.

### panHiTE accurately detects more intact TEs

By developing a novel LTR-RT detection algorithm and a method for recovering low-copy TEs, panHiTE can detect more intact TEs, many of which are absent from the gold-standard Repbase database. This highlights the potential of panHiTE as a valuable tool for complementing existing TE databases. As shown in Fig. 2a, a comparative analysis of pan-TE libraries generated by panHiTE and panEDTA across 32 *Arabidopsis* genomes, benchmarked against the Repbase database, reveals that the three databases share 235 TE families. Notably, panHiTE shares 323 TE families with Repbase that are not detected by panEDTA, whereas panEDTA shares only 32 TE families with Repbase that are absent in panHiTE. Further analysis of the 865 TEs uniquely identified by panHiTE reveals that 329 of these elements show significant overlap with known TE protein domains, suggesting their potential for autonomous mobility. The remaining 536 are regarded as non-autonomous (Fig. 2a). Among these, 141 belong to LTR terminals, 22 to LTR internals, 327 to TIRs, 44 to Helitrons, and 2 to non-LTRs. Within the 865 unique TEs discovered by panHiTE, the low-copy TEs recovered through the recovering low-copy TEs method are predominantly non-autonomous, including 22 TIRs and 4 Helitrons. Manual curation identified 1 Helitron and 4 TIRs as false positives, while the remaining 21 TEs were validated as genuine. These findings underscore the utility of panHiTE in expanding the catalog of known TEs.

**Fig. 2.**
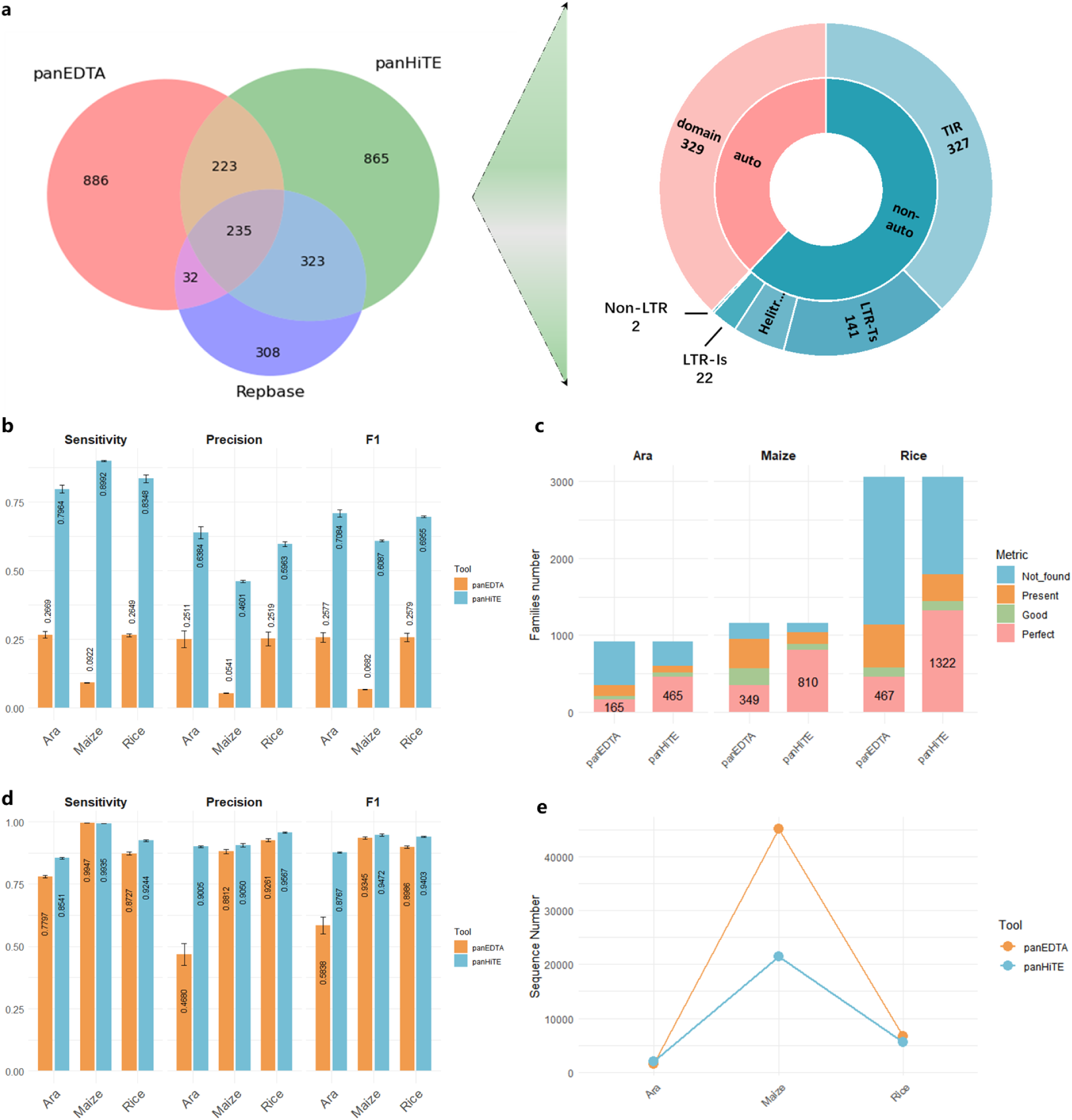
Analysis of pan-TE libraries. **a.** Comparison of pan-TE libraries generated by panHiTE and panEDTA across 32 *Arabidopsis* genomes, benchmarked against the Repbase database. **b, c, d.** Performance evaluation of panHiTE and panEDTA across 32 *Arabidopsis*, 26 maize, and 16 rice pan-genomes using three benchmarking methods: BM_HiTE (**b**), BM_RM2 (**c**), and BM_EDTA (**d**). **e.** Comparison of the number of sequences in the pan-TE libraries generated by panHiTE and panEDTA. Ara: Arabidopsis.

A robust and well-established evaluation framework is essential for accurately assessing the performance of computational methods in a competitive manner. In recent studies, three benchmarking methods introduced in the context of HiTE, RepeatModeler2, and EDTA have been demonstrated to provide more reasonable and comprehensive evaluations (referred to as BM_HiTE, BM_RM2, and BM_EDTA hereafter). BM_RM2 is based on RepeatMasker and a custom bash script provided by RepeatModeler2. An ideal TE library should consist exclusively of full-length TE models, which can be evaluated using the *Perfect* indicator from BM_RM2. TE models are classified as *Perfect* matches if they exhibit >95% sequence similarity, >95% length coverage, and <5% divergence relative to the family consensus in the gold standard library.

Both BM_HiTE and BM_EDTA involve comparing test annotations to curated genome annotations and calculating true positives (TP) and false negatives (FN). However, the key distinction lies in the criteria for defining true positives. BM_EDTA considers any sequence that has any length of match with the standard library as a true positive. In contrast, BM_HiTE requires the overlap between the test and curated TE sequences to exceed a stringent threshold (e.g., 95% of their respective lengths) to qualify as a true positive. All other alignments, including those with significant shifts, longer sequences containing the true TE, and fragmented TE sequences, are classified as false positives. This stringent criterion makes BM_HiTE more suitable for evaluating high-quality, full-length TE libraries by minimizing the influence of fragmented sequences on the results.

Given that Repbase contains high-quality TE consensus sequences for different subtypes within the same species, we selected it as the gold standard for evaluating pan-TE libraries. panHiTE demonstrates superior overall performance compared to panEDTA across all three benchmarking methods. Using BM_HiTE, panHiTE significantly outperforms panEDTA in sensitivity, precision, and F1-score. Specifically, panHiTE achieves 2.75-fold, 8.93-fold, and 2.70-fold improvements in F1-score based on pan-genomes of 32 *Arabidopsis*, 26 maize, and 16 rice accessions, respectively (Fig. 2b). In terms of the number of sequences in the pan-TE library, panHiTE and panEDTA generate comparable quantities for *Arabidopsis* and rice pan-genomes. However, for large genomes such as maize, panHiTE produces only half the number of sequences compared to panEDTA (Fig. 2e). Despite this, panHiTE generates 2–3 times more *Perfect* TE families than panEDTA. For example, using BM_RM2, panHiTE detects 465, 810, and 1,322 *Perfect* TE families in the pan-genomes of *Arabidopsis*, maize, and rice, respectively, while panEDTA detects only 165, 349, and 467, representing 2.82-fold, 2.32-fold, and 2.83-fold improvements (Fig. 2c).

As previously noted, BM_EDTA cannot distinguish between full-length TEs and short homologous false positives or fragmented sequences, potentially overestimating the performance of tools that produce fragmented TEs. Nevertheless, panHiTE still outperforms panEDTA on the pan-genomes of *Arabidopsis*, maize, and rice using BM_EDTA. For example, panHiTE attains F1-scores of 0.8767, 0.9472, and 0.9403, outperforming panEDTA, which achieves 0.5838, 0.9345, and 0.8986 (Fig. 2d). Notably, while panEDTA performs similarly to panHiTE on maize, its pan-TE library contains more than double the number of sequences, yet fewer than half the number of *Perfect* TE families. This indicates that library of panEDTA includes a significant proportion of fragmented TE families, which exhibit high homology to curated TE consensus sequences, allowing it to achieve high performance using BM_EDTA. However, when evaluated using BM_HiTE, the performance of panEDTA on maize drops significantly, with an F1-score of only 0.0682. In summary, the latest version of HiTE achieves substantial improvements through the development of the novel LTR-RT detection tool FiLTR and continuous enhancements in detecting other types of TEs. Building on this, panHiTE incorporates the upgraded HiTE along with a progressive redundancy removal algorithm and a low-copy TE recovery strategy, delivering consistently superior performance over panEDTA across nearly all pan-genomes. These advancements highlight its robustness and accuracy in pan-TE annotation.

### panHiTE accurately discovers the pan-TE landscape

Full-length TEs generally maintain intact structures and essential protein domains, suggesting they retain transposition activity^33,44^. In contrast, fragmented TEs, which have accumulated mutations or undergone significant structural variations leading to the loss of critical protein domains, often represent remnants of ancient TEs^45^. To facilitate our analysis, we categorized TEs into two groups: full-length TEs and complete TEs (including both full-length and fragmented TEs). We have identified 2,023 TE families from the full-length TEs, which are present in at least one of the 32 *A. thaliana* genomes. When considering the complete TEs, the number of TE families increased to 2,139. Among the full-length TEs, 564 (27.9%) are present in all 32 genomes and classified as core TE families, while 442 (21.8%) are present in 25–31 accessions (≥80% of the accessions) and defined as softcore, 895 (44.2%) are present in 2–24 genomes and classified as dispensable, and the remaining 122 (6%) are found in only one genome and designated as private TE families (Fig. 3a). For the complete TEs, 1,633 (76.3%) are core TE families, 212 (9.9%) are softcore, 268 (12.5%) are dispensable, and 26 (1.3%) are private.

**Fig. 3.**
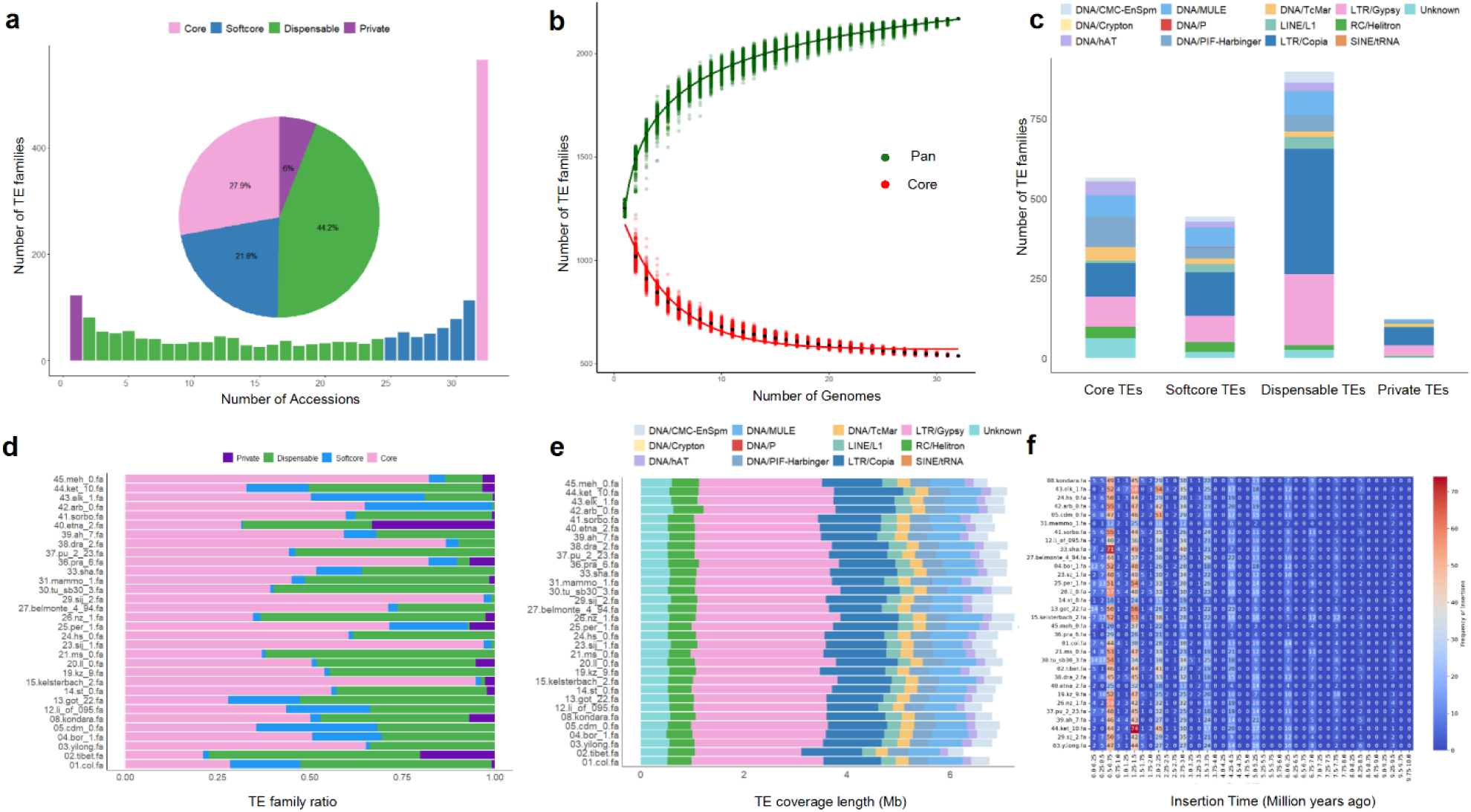
The full-length TE landscape of 32 *Arabidopsis thaliana* accessions. **a.** The number and proportion of core (red), softcore (blue), dispensable (green), and private (purple) TE families within the pan-genomes. **b.** A model depicting the number of TE families in the core and pan-genomes. Black dots indicate mean values. The number of pan-TE families grows with each additional accession, reaching a maximum of 2,023 TE families, with an extrapolated prediction of 2,383 ± 3 TE families. In contrast, the number of core-TE families declines with each added accession, stabilizing at 564 TE families, and is predicted to reach 594 ± 7 TE families. (± values represent the standard error). **c.** The composition of different TE superfamily types within the pan-TE library. **d, e.** The TE family ratio (**d**) and TE coverage length (**e**) across the genome assembly. **f.** The distribution of LTR-RT insertion times across all accessions.

For the full-length TEs, the number of pan-TE families increases as more accessions are added, eventually reaching a maximum of 2,023 TE families, with an extrapolated estimate of 2,383 ± 3 TE families. In contrast, the number of core-TE families declines with each additional accession, stabilizing at 564 TE families and predicted to reach 594 ± 7 TE families (Fig. 3b). The ± values represent the standard error. Similarly, for the complete TEs, the number of pan-TE families grows with each additional accession, reaching a maximum of 2,039 TE families, with an extrapolated prediction of 2,072 ± 0.3 TE families, while the number of core-TE families declines with each added accession, stabilizing at 1,593 TE families, and is predicted to reach 1,599 ± 4 TE families.

Among the complete TE families, DNA transposons and long terminal repeat (LTR) retrotransposons constituted the majority of TEs, with LTR/Copia being the most abundant (Fig. 3c). We observed significant variation in the proportions of core, softcore, dispensable, and private TE families across different genomes, indicating that the 32 *A. thaliana* accessions have undergone independent evolutionary processes (Fig. 3d). Additionally, the size differences among genomes are primarily attributed to variations in the coverage of LTR/Gypsy and LTR/Copia elements (Fig. 3e).

By analyzing the insertion times of LTR-RTs, we have observed periodic bursts of activity in both types of LTR-RTs. Specifically, since 500 million years ago, there has been a noticeable burst approximately every 500,000 years. This pattern is observed across all genomes, indicating that LTR-RT bursts are rhythmic events independent of the geographical distribution or varieties of the accessions (Fig. 3f).

### Population-specific TE insertions contributing to differential gene expression

By carrying cis-regulatory sequences, TEs can reshape gene regulatory networks through duplication and insertion, redistributing transcription factor (TF) binding sites and creating new enhancer activities^46,47^. TE insertions can disrupt regulatory or coding regions of target genes, thereby suppressing their expression^48^. They can also alter the splicing patterns of target genes, leading to the production of different transcripts or protein isoforms^49^. Large TE insertions can cause chromosomal rearrangements, such as deletions, duplications, or translocations, thereby affecting the expression of multiple genes^3^. To minimize the impact on host genes, long TE insertions such as LTR-RTs are often located far from host coding genes, resulting in an inverse distribution between LTR-RTs and coding genes^50^. At the same time, TE expression itself— through the transcript, the act of transcription, or subsequent TE replication intermediates—can regulate gene expression and chromatin accessibility^48,51^, activate cellular signaling pathways^52^ (e.g., interferon response or RNA interference responses), and trigger processes such as aging or antiviral activities^53^.

panHiTE leverages RNA-seq data to quantify TE and gene expression across populations, enabling the detection of population-specific TE insertions and analysis of their impact on differential gene expression, ultimately contributing to phenotypic differences among species. First, panHiTE quantifies TE expression levels. We have observed that certain accessions, including “Tibet-0”, “Bor-1”, “Sha”, “Etna-2”, and “Ket-10”, exhibit higher TE expression levels than others, suggesting increased TE activity in these accessions (Fig. 4a). Additionally, quantification at the superfamily level revealed that LTR/Copia, LTR/Gypsy, and DNA/TcMar show numerous outliers, with median expression levels close to zero (Fig. 4b). This suggests that while most copies of these superfamilies are genomic “fossils”, many still exhibit unusually high activity. Intriguingly, both active and remnant TEs are recognized as participants in evolutionary innovation and biological processes^45,54^, such as embryonic development^55^, as well as in human diseases and cancer^7,56^. We also observe that TEs from the same family may show significant expression variation among accessions (Fig. 4c). For example, “38-LTR_193_int” is highly expressed in “Tibet-0” but shows low expression in all other accessions. This TE expression specificity may be one of the key factors contributing to species divergence.

**Fig. 4.**
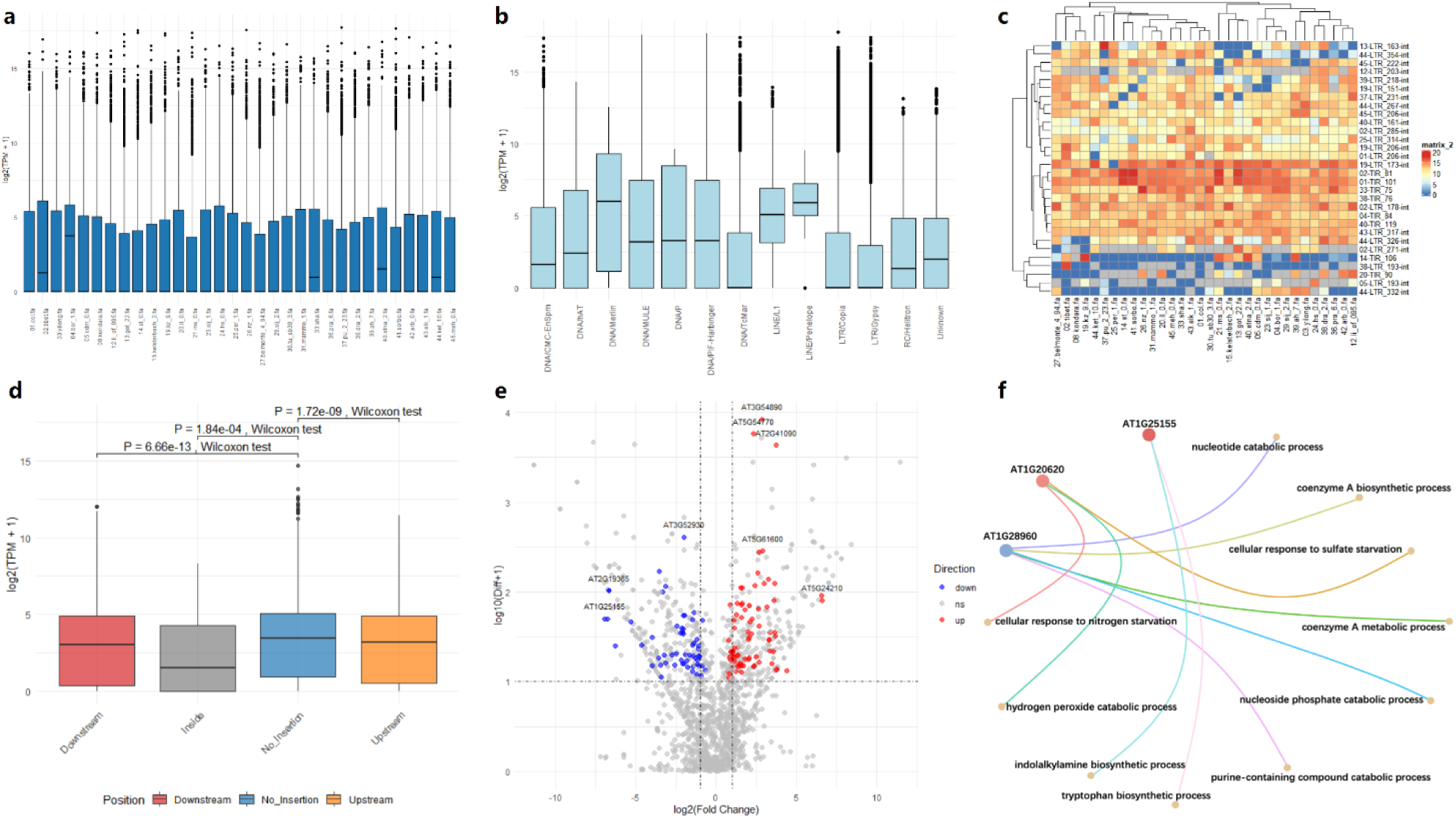
Analysis of TE and gene expression across 32 different *Arabidopsis* accessions. **a.** Distribution of TE expression levels across different accessions. **b.** Expression distribution of TEs across different superfamilies. **c.** Heatmap matrix showing TE activity differences across accessions. TEs from the same family may show significant expression variation among accessions. **d.** The impact of TE insertions on gene expression varies depending on their location relative to genes (downstream, internal, or upstream). Significance was determined using the Wilcoxon rank-sum test with Benjamini-Hochberg p-value adjustment to control for false discovery rate, and No_Insertion was used as the reference group. **e.** Volcano plot displaying candidate loci where TE insertions cause differential gene expression (referred to as “TE-induced differential expression loci” or TIDELs). Red dots represent genes with significantly upregulated expression due to TE insertions, blue dots represent downregulated genes, and gray dots indicate no significant difference or insufficient sample size for statistical support. **f.** Biological processes associated with three significant TIDELs. These genes play a role in the synthesis of important enzymes.

panHiTE can perform TE-gene positional association analysis (Fig. 1b). We compared the expression levels of genes with TE insertions upstream, internally, downstream, and those without TE insertions. We have observed that genes with internal TE insertions in *Arabidopsis thaliana* Col-0 exhibit significantly lower expression levels than those without (Fig. 4d), suggesting that TE insertions within genes often disrupt their normal expression. Similar patterns are observed across all other 31 ecotypes.

Generally, it is believed that TEs influence species traits by regulating gene expression. To analyze the causes of trait-specific variations within populations, we use loci as analytical units, dividing samples into those without TE insertions and those with TE insertions (upstream, internal, or downstream). We then calculate gene expression differences between these groups to identify candidate loci where TE insertions cause differential gene expression (referred to as “TE-induced differential expression loci” or TIDELs). These loci provide a foundation for studying population evolution and species improvement. In 32 *Arabidopsis* accessions, we have identified hundreds of TIDELs (Fig. 4e), which are associated with various biological processes (Fig. 4f) and are involved in the synthesis of important enzymes.

### FiLTR accurately detects LTR-RTs

Long terminal repeat retrotransposons (LTR-RTs) are abundant in both plant and animal genomes, playing key roles in maintaining genome stability and regulating gene expression^57,58^. Accurate identification of LTR-RTs is vital for high-quality genome annotation. However, existing methods suffer from the lack of accuracy and robustness, leading to a high rate of false positives. Here, we implement FiLTR, a novel approach that combines deep learning with boundary detection algorithms to enhance LTR-RT detection. FiLTR begins by screening candidate LTR-RTs to remove those with contaminated internals and terminals composed of tandem repeats (Fig. 5a), and refines the boundaries of the candidates (Fig. 5b). Next, FiLTR constructs a flanking matrix for LTR-RTs, uncovering a pattern of half-homology and half-randomness in the flanking regions (Fig. 5c). Based on this discovery, FiLTR encodes the bases of the flanking matrix with discrete color values, extracts multi-dimensional features, and applies a deep learning model designed for high-sensitivity detection of authentic LTR-RTs (Fig. 5d). Subsequently, FiLTR implements a homology-based boundary detection algorithm to accurately identify LTR terminal boundaries and filter out false positives overlooked by the deep learning model. For more information on the filtering algorithm based on the flanking matrix of FiLTR, please refer to the Methods section. Additionally, FiLTR identifies false positives composed of other TEs such as terminal inverted repeats (TIRs), Helitrons, and non-LTRs (Fig. 5e). After this, the full-length LTR-RTs are aligned with the genome to identify their intact copies. For single-copy LTR-RTs, their authenticity is determined by checking for the presence of intact LTR domains and target site duplications (TSDs), as shown in Fig. 5f. Finally, a progressive redundancy removal algorithm is developed to generates LTR consensus sequences while reducing fragmentation and nested LTR-RTs (Fig. 5g).

**Fig. 5.**
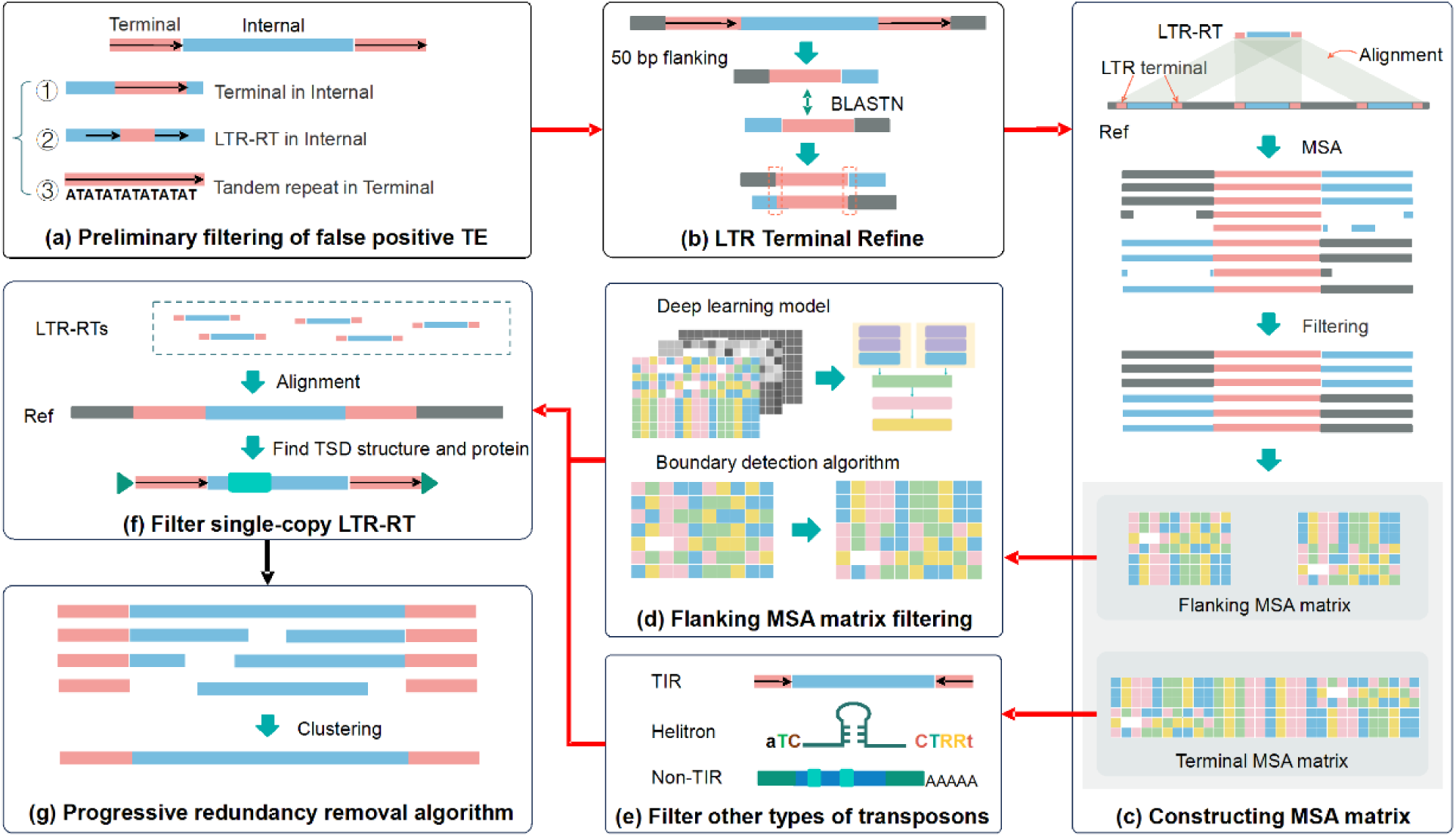
Workflow of the FiLTR. **a.** Preliminary screening for false positives. **b.** Correction of LTR boundaries. **c.** Construction of flanking matrix for LTR terminal, with colored blocks representing different bases. **d.** Identification of false positives using a flanking matrix filtering method. **e.** Filtering false-positive terminals consisted of other TEs. **f.** Filtering method for single-copy LTR-RTs. **g.** Generation of LTR library using a progressive redundancy removal algorithm.

We refer to tools such as LTR_harvest, LTR_finder, and LtrDetector as primary LTR-RT detection tools. As shown in Figs. 6a and b, they exhibit high sensitivity but also produce a large number of false positives. For example, in *Drosophila melanogaster*, LtrDetector consistently exhibits high sensitivity but low precision, achieving a sensitivity of 0.8707 and a precision of 0.0068 with BM_HiTE, and a sensitivity of 0.9945 and a precision of 0.3675 with BM_EDTA (Fig. 6). Similarly, in *Oryza sativa*, LtrDetector attains a sensitivity of 0.943 with a precision of 0.0419 using BM_HiTE, and a sensitivity of 0.9905 with a precision of only 0.4397 using BM_EDTA (Fig. 6). LTR_harvest and LTR_finder produce similar results. Tools such as LTR_Retriever, Inpactor2, and FiLTR serve as advanced LTR-RT detection tools. LTR_Retriever implements rule-based filters, which significantly improve precision compared to tools like LTR_harvest, LTR_finder, and LtrDetector, but there remains room for further improvement. Inpactor2, a deep learning tool based on LTR_finder, performs comparably to LTR_Retriever in *Oryza sativa* and *Danio rerio* but underperforms in *Drosophila melanogaster*, likely due to its training on specific species.

**Fig. 6.**
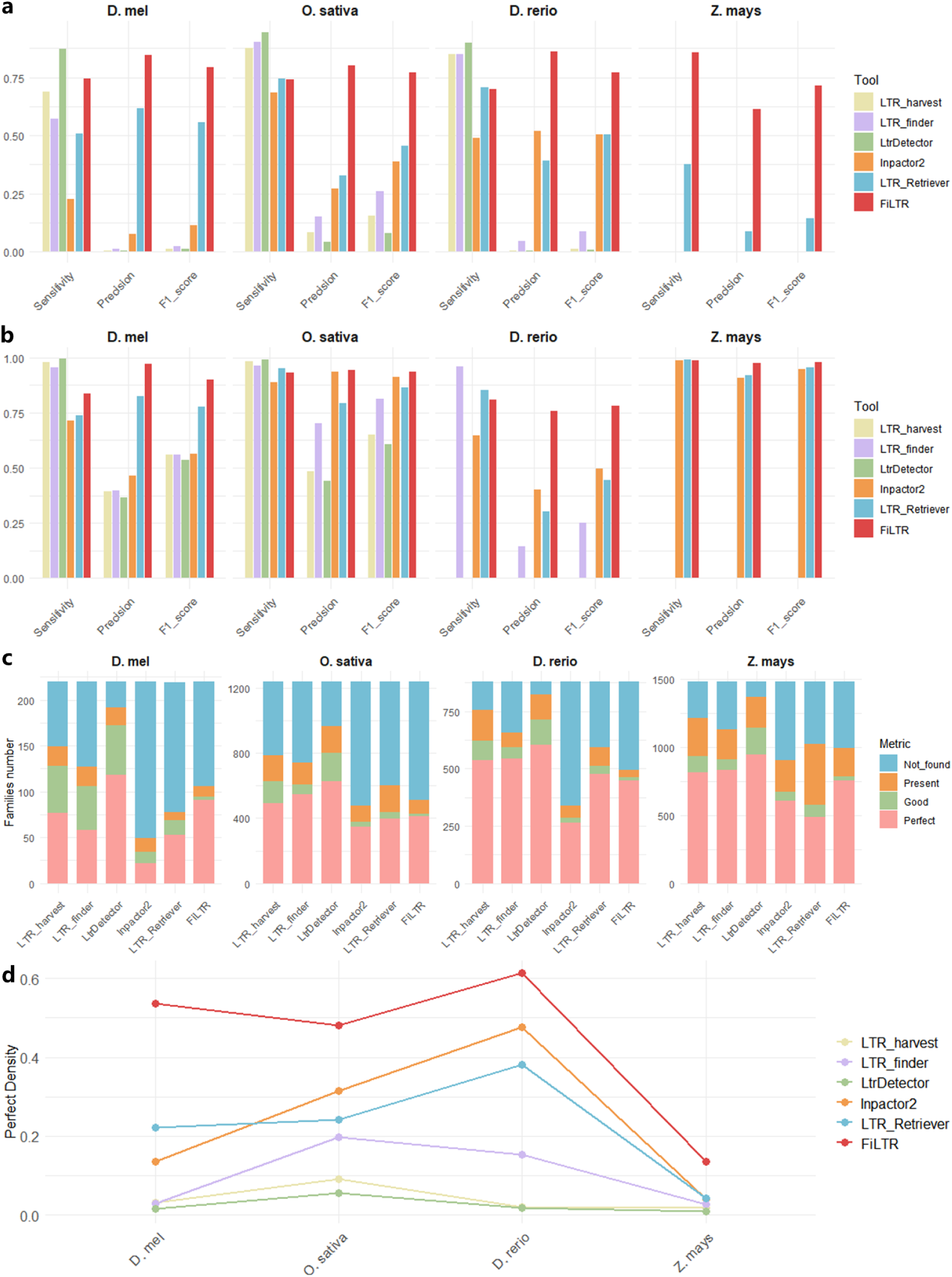
Performance evaluation of different tools using BM_HiTE, BM_EDTA, and BM_RM2. **a, b.** The bar chart illustrates performance evaluated by the BM_HiTE (**a**) and BM_EDTA (**b**) methods. **c.** The stacked bar chart shows performance assessed by the BM_RM2 method. **d.** The line graph displays the performance based on the *Perfect Density* metric. A higher *Perfect Density* indicates that the tested TE library contains fewer redundant, fragmented, or false-positive sequences. Note that some tools produced excessively large results in *D. rerio* or *Z. mays*, making their assessment impossible; these cases are represented by zeros. Both FiLTR and LTR_Retriever utilize outputs from LtrDetector as input.

Compared to competing tools, FiLTR improves the detection performance of LTR-RTs by filtering out many false positives, showing enhancements with both BM_HiTE and BM_EDTA. For example, FiLTR improves the F1 score of LtrDetector from 0.0803 to 0.757 in *Oryza sativa* using BM_HiTE, representing a substantial enhancement. This corresponds to an approximately 66% improvement over LTR_Retriever, which achieves an F1 score of 0.4555 (Fig. 6a). When using BM_EDTA, FiLTR further increases the F1 score of LtrDetector from 0.6090 to 0.9333 in *Oryza sativ*a, exceeding the F1 score of 0.8657 obtained by LTR_Retriever (Fig. 6b). Moreover, FiLTR demonstrates higher F1 scores across other species (Fig. 6). Notably, in large genomes such as *Danio rerio* and *Zea mays*, the large LTR libraries generated by Inpactor2, LTR_harvest, LTR_finder, and LtrDetector present challenges for evaluation using BM_HiTE and BM_EDTA (Figs. 6a and b).

BM_RM2 evaluates the quality of TE libraries based on the presence of full-length TE models. Notably, aside from the *Perfect* metric, all other metrics (*Good* and *Present*) indicate some degree of fragmentation. Primary LTR-RT detection tools, such as LTR_harvest, LTR_finder, and LtrDetector, generate a high number of *Perfect* models while also producing a substantial proportion of *Good* and *Present* models (Fig. 6c). Compared to these primary detection tools, FiLTR reduces the number of *Perfect* models but also decreases the counts of *Good* and *Present* models. When comparing advanced LTR-RT detection tools, LTR_Retriever outperforms Inpactor2 in detecting *Perfect* models for most species but also results in a higher count of *Present* models. FiLTR outperforms LTR_Retriever in most species, with the exception of *Danio rerio*. While FiLTR shows slightly fewer *Perfect* models for *Danio rerio*, it also generates fewer *Good* and *Present* models compared to LTR_Retriever. It is crucial to highlight that an ideal TE library should aim to maximize the number of *Perfect* models while minimizing *Good* and *Present* models^34^.

Ideally, a TE library should align perfectly with the family counts of a curated TE library. However, due to the repetitive nature of TEs, automated tools often inevitably produce significant redundancy, fragmentation, and false positives, frequently requiring extensive manual curation and editing. To address this issue, we introduce a new metric, *Perfect Density*, which quantifies the proportion of *Perfect* LTR-RT models in the library (Methods). A *Perfect Density* value closer to 1 indicates lower redundancy and fragmentation in the TE library. As shown in Fig. 6d, primary LTR-RT detection tools such as LTR_harvest, LTR_finder, and LtrDetector perform poorly on the *Perfect Density* metric, highlighting issues with redundancy, false positives, and fragmentation in their LTR libraries. In contrast, advanced LTR-RT detection tools like Inpactor2, LTR_Retriever, and FiLTR significantly improve the *Perfect Density* metric. Notably, FiLTR achieves the highest *Perfect Density* across all species, producing accurate and compact libraries.

## Discussion

Advancements in sequencing technologies have enabled the analysis of an increasing number of individuals, making population-specific TE detection a powerful tool for studying species evolution and improving crop breeding^59,60^. However, existing tools for fast and accurate pan-genome TE detection remain limited, especially for large and complex genomes. In this study, we develop panHiTE, a Nextflow-based workflow designed for population-scale TE detection and annotation using pan-genomes. To enhance the accuracy of LTR-RT detection, we develop FiLTR as a replacement for the previously used LTR_Retriever in HiTE. FiLTR integrates deep learning with boundary detection algorithms, significantly improving the completeness and accuracy of LTR identification compared to existing methods. Additionally, panHiTE leverages pan-genomes to recover low-copy TEs missed in single-genome analyses, outperforming panEDTA in both sensitivity and precision of TE detection.

panHiTE provides a comprehensive TE landscape across pan-genomes, capturing both full-length and fragmented TEs. It categorizes TE families into core, softcore, dispensable, and private groups (Fig. 3a) and models the accumulation of TE families using saturation curves (Fig. 3b). Additionally, it characterizes the composition of TE superfamilies (Fig. 3c), calculates TE family ratios (Fig. 3d), evaluates TE coverage across genome assemblies (Fig. 3e), and estimates the distribution of LTR-RT insertion times across accessions (Fig. 3f). Our analysis of 32 *Arabidopsis* ecotypes revealed periodic bursts of LTR insertions occurring approximately every 500,000 years.

By integrating RNA-seq data, panHiTE quantifies TE and gene expression across populations, facilitating the identification of population-specific TE insertions and their impact on differential gene expression. This capability provides insights into phenotypic variation among species. For instance, we have observed that certain accessions exhibit higher TE expression levels, indicating increased TE activity (Fig. 4a). While many TEs are genomic “fossils”, some superfamilies remain highly active. Notably, TEs from the same family can display significant expression variation across accessions (Fig. 4c), underscoring their potential role in species divergence. panHiTE also performs TE-gene positional association analysis (Fig. 1b). By comparing gene expression levels in the presence of TE insertions located upstream, internally, downstream, or absent, we have found that genes with internal TE insertions exhibit significantly reduced expression (Fig. 4d). This suggests that TE insertions within genes often disrupt normal gene function. To further investigate the functional impact of TEs, we identified TE-induced differential expression loci (TIDELs) by comparing gene expression differences between groups with and without TE insertions. These loci represent valuable candidates for studying population evolution and species improvement. In 32 *Arabidopsis* accessions, we have identified hundreds of TIDELs (Fig. 4e) linked to various biological processes (Fig. 4f) and involved in the synthesis of essential enzymes.

In summary, by implementing a more precise LTR-RT detection algorithm and continuous improvements to HiTE, panHiTE offers the most accurate pan-TE detection and annotation method currently available. Built on Nextflow, panHiTE leverages high-performance computing platforms to rapidly perform TE detection and annotation on large-scale population genomes. panHiTE has been successfully applied to large-scale pan-genome analyses, including hundreds of *Brassica* genomes and 17 ultra-large *Triticum* genomes. We anticipate that the methods proposed in this study will contribute to population-scale variation research, with potential applications in areas such as human disease and crop breeding.

## Methods

### Recovering low-copy TEs using pan-genomes

The accurate identification of low-copy TEs remains challenging, primarily due to ambiguities in defining their precise boundaries and structural characteristics. These limitations frequently lead to false-positive annotations or even the complete omission of low-copy TEs during detection processes. Pan-genomes provide a method to address this issue by enabling cross-genome realignment of low-copy TEs. This approach aggregates sufficient homologous copies to validate their authenticity. In panHiTE, sequences initially classified as low-copy candidates meet two criteria: (1) fewer than five full-length copies (based on an empirical threshold) and (2) absence of canonical TE features such as terminal inverted repeats (TIRs) or conserved TE domains. These candidates are realigned across multiple genomes, followed by collection of their aligned copies and flanking sequences. Finally, multiple sequence alignment is performed to assess whether these copies exhibit bona fide TE boundaries and structural integrity.

### Progressive redundancy removal algorithm

Extensive insertions and deletions within TEs cause variability among instances of the same TE family, resulting in numerous redundant and fragmented sequences. To address this issue, we have developed a progressive redundancy removal algorithm. First, a rapid fault-tolerant alignment clustering algorithm^40^ is designed to align sequences with substantial insertions and deletions, allowing for the quick clustering of divergent instances from the same TE family. Next, multiple sequence alignments are performed for each cluster, and the Ninja algorithm^61^ is applied for finer clustering. Finally, consensus sequences are generated based on the majority rule, effectively reducing redundancy and improving the accuracy of TE family models.

### TE and gene quantification

We began by preprocessing the raw RNA-seq reads to remove adapter sequences and filter out low-quality reads. For single-end reads, we used Trimmomatic^62^ with the following parameters: SE ILLUMINACLIP:TruSeq3-SE.fa:2:30:10 LEADING:3 TRAILING:3 SLIDINGWINDOW:4:15 MINLEN:36 TOPHRED33. For paired-end reads, we applied the parameters: PE ILLUMINACLIP:TruSeq3-PE.fa:2:30:10 LEADING:3 TRAILING:3 SLIDINGWINDOW:4:15 MINLEN:36 TOPHRED33. The resulting high-quality reads are then aligned to the reference genomes using HISAT2^63^. To quantify expression levels, we calculated transcripts per million (TPM)^64^ for both TEs and genes using featureCounts^65^. Multi-mapping reads are excluded from TE quantification using featureCounts (countMultiMappingReads=FALSE) to ensure accuracy.

### Identification of gene differential expression induced by TE insertions

We begin by analyzing the full-length TE annotations and gene annotation files to identify TE insertions within a 10 kb flanking region (upstream or downstream) of each gene. For each gene in the population, we categorized it into four groups based on the presence or absence of TE insertions: (1) upstream insertions, (2) internal insertions, (3) downstream insertions, and (4) no insertions (control group). Subsequently, we compared the gene expression levels between the control group (no TE insertions) and each of the other three groups (with TE insertions) to identify genes exhibiting significant differential expression. We established two parameters, *min_sample_threshold* and *min_diff_threshold*, to identify genes with significant differential expression. The *min_sample_threshold* defines the minimum sample size required for a valid comparison between two groups. For instance, when comparing the internal insertions group with the no insertions group, a *min_sample_threshold* of 5 indicates that at least one of the two groups must contain more than 5 samples; otherwise, the comparison is considered statistically unreliable. The *min_diff_threshold* specifies the minimum absolute difference between the lowest value in one group and the highest value in the other group. Only when the expression values between the two groups are sufficiently distinct is the TE insertion deemed to have a significant impact on gene expression levels.

### De novo LTR-RT detection

LtrDetector identifies potential repetitive sequences by calculating k-mer distance scores, allowing for the detection of a comprehensive set of candidates with long terminal repeats. It demonstrates superior sensitivity compared to LTR_harvest and LTR_finder, making it the preferred primary LTR-RT detection tool for FiLTR. To enable rapid detection of LTR-RTs in large genomes, the genome is divided into 10 Mb segments, and a highly parallelized version of LtrDetector is implemented to identify candidate LTR-RTs.

### Preliminary screening for false positives

LTR-RTs exhibit long terminal repeats, allowing structure-based detection tools such as LTR_harvest, LTR_finder, and LtrDetector to identify the potential LTR-RTs with high sensitivity. However, these tools often yield a significant number of false positives. To address this issue, we implement several screening steps for candidate LTR-RTs as follows:

i. Removal of LTR-RTs with internals containing LTRs. Given that the long terminal repeats of LTR-RTs exhibit a high degree of similarity, they can lead to recombination within the element^66,67^, resulting in complex LTR structures that can hinder accurate identification of full-length LTR-RTs. To ensure the detection of clean full-length LTR-RTs, we filter out candidate LTR-RTs that have their terminals located within their internal regions.
ii. Exclusion of candidates containing intact LTR-RT. We further remove candidates that contain intact LTR-RTs within their internal regions. These nested LTR-RTs can interfere with the generation of high-quality full-length LTR-RT libraries. Additionally, these inserted elements typically have intact LTR-RT copies located elsewhere in the genome, so filtering them out generally does not affect the overall performance of LTR-RT detection.
iii. Removal of LTR-RTs with terminals made up of tandem repeats. Tandem repeats are typically composed of simple repetitive units, and adjacent tandem repeats can easily form structures resembling long terminal repeats, potentially leading to false positive detections. To mitigate this issue, we use the TandemRepeatFinder^68^ to identify whether there are substantial amounts of tandem repeats present in the long terminal repeats and filter out false positive terminals made up of tandem repeats.

### Refinement of LTR terminal boundaries

LTR-RTs generated by structure-based detection tools may exhibit inaccuracies in their terminal boundaries. However, the filtering methods of FiLTR depend on the precise determination of LTR boundaries, requiring a refinement of these boundaries. In this step, we correct the LTR boundaries using the following approach: we first obtain the 5’- and 3’-LTR terminals and then extend each terminal sequence by 50 bp on both sides. Next, we perform a BLASTN^69^ alignment on the extended 5’- and 3’-LTR terminals and adjust the terminal boundaries based on the alignment results.

### Construction of flanking matrix for LTR terminals

To eliminate false positives, we construct a multiple sequence alignment (MSA) matrix based on the LTR terminal and its flanking regions. The process consists of three steps, illustrated with an LTR-RT example. First, the 5’-LTR terminal is aligned to the genome to identify its copies. Second, each copy is extended by 100 base pairs on both sides to capture flanking regions, and a multiple sequence alignment is performed to construct a flanking matrix. Finally, columns with a high proportion of gaps are removed based on a majority rule, preserving information density while minimizing the impact of sparse data.

### Filtering algorithm based on flanking matrix

When conducting multiple sequence alignment on all 5’-LTRs of the same family, the left flanking region appears random, while the right flanking region shows homology. This pattern arises from the stochastic nature of LTR-RT insertions and the inherent similarities within the internal regions of the same LTR-RT family. Similarly, for 3’-LTRs, the left flanking region displays homology, and the right flanking region exhibits randomness. Due to the high similarity between LTRs, copies derived from the 5’-LTR can originate from both the 5’ and 3’ ends. Assuming that 5’- and 3’-LTRs occur with equal frequency in the genome, the multiple sequence alignment of 5’-LTR copies reveals a pattern of half-homology and half-randomness. Consequently, when the flanking matrix is divided into the left flanking region, the central region, and the right flanking region, the central region of genuine LTR-RTs is marked by high homology, whereas the flanking regions exhibit a half-homology, half- randomness pattern.

### The deep learning model

To filter out false positives while retaining genuine LTR-RTs, we encode the bases of the flanking matrix with discrete color values, extract multi-dimensional features, and apply a deep learning model designed for high-sensitivity detection of authentic LTR-RTs.

i. Dataset construction. To construct the dataset, we have downloaded genome assembly data for over 1,700 species from the NCBI database and aligned the LTR-RTs from Repbase to these genomes to obtain their copies, ultimately constructing a multiple sequence alignment (MSA) matrix for the LTR flanking regions. As a result, we generate 9,354 positive samples across 238 species. Simultaneously, we utilize other TEs, including non-LTR, TIR, and Helitron elements from Repbase, segmenting them into three parts—head, middle, and tail sequences—and construct their corresponding flanking matrices. This process results in a total of 10,628 negative samples.
ii. Model architecture. The architecture of this neural network is built upon the ResNet^70^ model, incorporating three BasicBlock layers for feature extraction, which include bases, blocks, and supports. The bases features are generated by mapping the bases in the flanking matrix to discrete, color-coded values. For the blocks features, gaps in the flanking matrix are encoded as 0, while non-gap positions are set to 100, aiming to reduce noise interference for the model. The supports features are derived by calculating the proportion of each base within each column of the flanking matrix, enabling the model to differentiate between homologous and random columns. Central to ResNet architecture is the BasicBlock, which encapsulates the essence of residual connections. This block is designed to learn the residual function *F*(*x*) = *H*(*x*) − *x*, where *H*(*x*) is the desired mapping and *x* is the input. The output of the BasicBlock is formulated as:

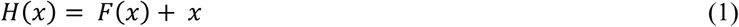 The residual connection is pivotal as it allows the input gradient to flow through directly, thus preventing the vanishing gradient problem. Here, *F*(*x*) is the composite function representing the sequence of convolutional and activation layers:

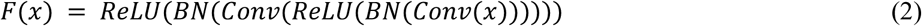 The inclusion of batch normalization after each convolutional layer standardizes the activations, which aids in faster training and enhanced generalization:

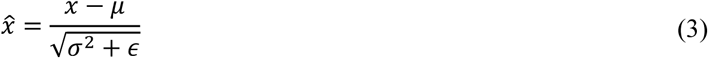 To further control the complexity of model, we add a global average pooling layer between the BasicBlock and the fully connected layer, which helps reduce the risk of overfitting and effectively decreases the number of model parameters and computations.
iii. Model training. During training, we set different random seeds to perform five independent random splits of the original dataset, with each training round comprising 25 epochs. The model parameters that yield the best F1 score on the test set are saved at the end of each round. Given that different TEs exhibit varying copy numbers in genomes, we aim to enhance the generalization of the model performance across TEs with different copy numbers. Therefore, we stratify the total training samples based on their copy numbers and randomly split each stratum in a 4:1 ratio, ultimately merging them into a training dataset and a validation dataset. We minimize the cross-entropy loss to guide the model in learning parameters on the training dataset, using a batch size of 256 and an initial learning rate of 0.001. The Adam optimizer controls the learning rate, and L2 regularization is applied by adding a penalty term to the loss function proportional to the square of the parameters, which helps prevent model overfitting.

### The boundary detection algorithm

The performance of the deep learning model depends on the diversity of patterns in the training data. Although the deep learning model can identify and exclude most false positives, some may still be overlooked. To address this issue, we design a homology-based boundary detection algorithm to further eliminate false positives not identified by the deep learning model. In the algorithm, we calculate the proportion of a specific nucleotide (A, T, C, G) in each column and take the maximum proportion as the homologous rate (*HR*) for that column. A column is considered valid if the homologous rate exceeds 50%, and invalid if it falls below this threshold.

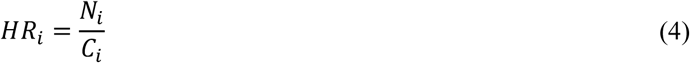

where *HR*_*i*_ is the homologous rate for column *i*, *N*_*i*_ is the count of the most abundant nucleotide (A, T, C, or G) in that column, and *C*_*i*_ is the total number of bases in that column.

We then use a sliding window to scan the valid columns in the flanking regions and calculate the average homology rate (*AHR*_*w*_) within the window. If the *AHR*_*w*_ surpasses a predetermined threshold (e.g., 85%), the window is classified as homologous; otherwise, it is considered non-homologous.

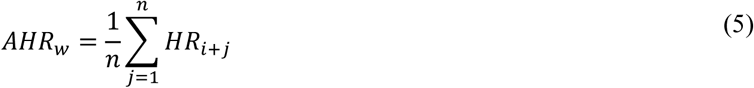

where *AHR*_*w*_ is the average homology rate within the window, *n* is the window size (e.g., 20 bp), and *HR*_*i*+*j*_ is the homologous rate for each valid column within the window.

For the left region of the flanking matrix, the window slides from the end toward the beginning to identify the first homologous column within the first non-homologous column as the left boundary (*LB*) of the matrix. For the right region, the window slides from the beginning toward the end to identify the last homologous column within the first non-homologous window as the right boundary (*RB*) of the matrix. Since all candidate LTR-RTs undergo boundary refinement (Fig. 5b), those whose identified boundaries differ significantly from the original boundaries are considered false positives and filtered out. Otherwise, the candidate LTR-RTs are considered true positives and retained.

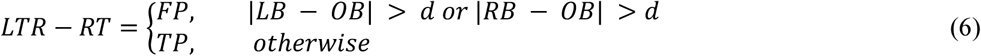

where *OB* represents the original boundaries, *LB* the identified left boundaries, and *RB* the identified right boundaries of the flanking matrix, and *d* the distance threshold between *LB*/*RB* and *OB*.

### Filtering terminals made up of other TEs

False positive LTR-RTs often result from other TEs that can form long terminal repeats. For example, two copies of the same DNA transposon can produce nearly identical long terminal repeats, resulting in a false positive. To address this issue, we implement a filtering approach based on the flanking matrix to filter out false positives generated by TIR (Terminal Inverted Repeat), Helitron, and Non-LTR transposons (Fig. 5e).

To filter TIR transposons, we first use a homology-based boundary detection algorithm^40^ to identify precise boundaries, then search for 2-11 bp TSDs adjacent to these boundaries. If copies with TSDs exceed 25% of the total or a threshold like 10, we check for TIRs in the consensus sequences. Consensus sequences with TIRs and TSDs are retained as reliable TIR transposons and used to exclude candidate LTR-RTs with similar terminals. A similar approach is applied for Helitron and Non-LTR transposons, using different methods to detect their structural features. For Helitrons, we look for 5’-TC…CTRR-3’ sequences, insertion AT sites, and hairpin structures, while for Non-LTRs, we focus on polyA tails and 8-20 bp TSDs. This approach effectively filters out false-positive terminals from TIR, Helitron, and Non-LTR transposons.

### Filtering method for single-copy LTR-RTs

Most candidate single-copy LTR-RTs are false positives, composed of other TEs. Given that these TEs typically exhibit high copy numbers, they can become widely distributed throughout the genome, significantly affecting the accuracy of annotation. To resolve this issue, we first align the candidate full-length LTR-RTs to the genome to identify single-copy LTR-RTs. We then compare these single-copy LTR-RTs against an LTR domain database and search for their TSDs. Only those single-copy LTR-RTs with both intact LTR domains and TSDs are considered true LTR-RTs.

## Data availability

The genome assemblies of 32 *Arabidopsis thaliana* ecotypes are available from Figshare (https://doi.org/10.6084/m9.figshare.21673895). The corresponding RNA-seq datasets can be accessed from the NCBI Sequence Read Archive (SRA) under BioProject accessions PRJNA187928, PRJEB15161, and PRJNA319904. Genome assemblies for 13 wild rice accessions are available at Figshare (https://figshare.com/s/32d0ac68cee7f647e1e3), while assemblies for 16 rice accessions can be accessed at the Rice Population Reference Panel (https://yongzhou2019.github.io/Rice-Population-Reference-Panel/data/). Genome assemblies of 26 *Zea mays* (maize) accessions are available from MaizeGDB (https://maizegdb.org/NAM_project). The genome assemblies of 17 wheat accessions are accessible via the National Genomics Data Center (NGDC) BioProject PRJCA021345 (https://ngdc.cncb.ac.cn/bioproject/browse/PRJCA021345). Reference genomes for *Drosophila melanogaster* (*Release 6 plus ISO1 MT*), *Oryza sativa* (*IRGSP-1.0*), *Danio rerio* (*GRCz11*), *Zea mays* (*Zm-B73-REFERENCE-NAM-5.0*), and *Arabidopsis thaliana* (*TAIR10.1*) are available from NCBI GenBank (https://www.ncbi.nlm.nih.gov/genome/). The curated TE libraries used in this study are available through a paid subscription to Repbase (https://www.girinst.org/repbase/). All TE and pan-TE libraries generated in this study are publicly available in the GitHub repository (https://github.com/CSU-KangHu/panTE_annotation) and Zenodo.

## Code availability

panHiTE is publicly accessible on GitHub [https://github.com/CSU-KangHu/HiTE] and Zenodo.

